# Dietary sphinganine is selectively assimilated by members of the gut microbiome

**DOI:** 10.1101/2020.06.08.140665

**Authors:** Min-Ting Lee, Henry H. Le, Elizabeth L. Johnson

**Author notes:** Abbreviations: DHCeramide, dihydroceramide; FACS, Fluorescence-activated cell sorting; FSC, forward scatter; HMO, human milk oligosaccharides; LC-HRMS, liquid chromatography coupled to high-resolution mass spectrometry; NSP, non-sphingolipid-producing; PAA, palmitic acid alkyne; PFA, paraformaldehyde; SAA, omega-alkynyl sphinganine; 16S sequencing, 16S rRNA gene sequencing; SP, sphingolipid-producing; SPT, serine palmitoyltransferase; SSC, side scatter.

## Abstract

Functions of the gut microbiome have a growing number of implications for host metabolic health, with diet being one of the most significant influences on microbiome composition. Compelling links between diet and the gut microbiome suggest key roles for various macronutrients, including lipids, yet how individual classes of dietary lipids interact with the microbiome remain largely unknown. A class of lipids known as sphingolipids are bioactive components of most foods and are produced by prominent gut microbes. This makes sphingolipids intriguing candidates for shaping diet-microbiome interactions. Here, we use a click-chemistry based approach to track the incorporation of bioorthogonal dietary omega-alkynyl sphinganine (sphinganine alkyne – SAA) into the gut microbial community (Click). Identification of microbe and SAA-specific metabolic products was achieved by fluorescence-based sorting of SAA containing microbes (Sort), 16S rRNA gene sequencing to identify the sphingolipid-interacting microbes (Seq), and comparative metabolomics to identify products of SAA assimilation by the microbiome (Spec). Together this approach, Click-Sort-Seq-Spec (ClickSSS), revealed that SAA-assimilation was nearly exclusively performed by gut *Bacteroides*, indicating that sphingolipid-producing bacteria play a major role in processing dietary sphinganine. Comparative metabolomics of cecal microbiota from SAA-treated mice showed conversion of SAA to a suite of dihydroceramides, consistent with metabolic activity via *Bacteroides* and *Bifidobacterium*. Additionally, other sphingolipid-interacting microbes were identified with a focus on an uncharacterized ability of *Bacteroides* and *Bifidobacterium* to metabolize dietary sphingolipids. Therefore, ClickSSS provides a platform to study the flux of virtually any alkyne-labeled metabolite in diet-microbiome interactions.

## Introduction

The mammalian intestinal microbiome is a complex and dynamic unit that influences host metabolism (1–3) and is significantly impacted by environmental factors (4, 5). There are many deterministic factors that shape the gut microbial consortium, yet diet has consistently been regarded as a dominant driver (6–8). In mammals, the intimate relationship between diet and host microbiome is particularly notable during the postnatal period where mother’s milk greatly functions to support the development of healthy symbiosis between infants and their developing gut microbial communities (9–13). This can occur when breastfed infants are exposed to components of human milk that are accessible nutrient sources for beneficial microbes. For example, indigestible carbohydrates in human milk and their impact on the gut microbiome have been extensively studied in this regard (14–19). There are other macronutrient components of human milk with uncharacterized functions in nutrient-gut microbiome interactions that have the potential to influence the development of the gut microbiome. Bioactive lipids in mammalian diets are a diverse suite of metabolites that serve essential functions in host metabolic health and microbial community composition (20). Changes in the fat composition of diets can have significant effects on gut microbial communities (21) yet little is known about which specific lipid structures interact with microbes in the gut and how specific species are able to uptake and metabolize dietary lipids. Classes of bioactive lipids that are metabolically accessible to the microbiome could serve as mechanisms to exert host control over gut microbial composition.

Sphingolipids are a class of bioactive lipids that are large components of the milk fat globule membrane surrounding the triglyceride fraction of human milk (22, 23). Bioactive sphingolipids in human milk have been shown to influence microbial growth *in vitro* (24–26) but not much is known on how these nutrients maybe utilized by the gut microbiome *in vivo*. Dietary sphingolipids vary in their structural composition, but the unifying characteristic is their sphingoid base, the initiating building block from which further modifications can introduce vast structural heterogeneity giving rise to more complex sphingolipids (27–29). Before being absorbed, digestive processes hydrolyze complex dietary sphingolipids, liberating the constituent components, including the sphingoid base (27–32). Although it exists as one of the major sphingoid bases in human milk (33), the interactions of dietary sphinganine (SA) and the gut microbiome remain largely unexplored. Prominent microbes of the infant gut have the capacity to synthesize sphingolipids (34) and the presence of microbiome-derived sphingolipids have been shown to be critical for proper immune system conditioning (35, 36). Given that sphingolipids could enhance the fitness of these beneficial microbes, it is important to understand if and how dietary sphingolipids interact with the microbiome.

As lipids are ubiquitous in nature, determining their origins and fates becomes a major challenge in studying both dietary lipids, and lipids in general. As sphingolipids can be derived from the host, the microbiome, and diets, specialized techniques are necessary to determine the origin of unique sphingolipid signals. Conventional efforts to monitor host uptake of dietary lipids employ radiolabeled or isotope-modified probes (27, 37–39) which require special accommodations and offer limited downstream application, prompting us to seek practical alternatives. To efficiently track the fate of dietary sphingolipids and identify sphingolipid-interacting microbial signatures, we leveraged a burgeoning tool — bioorthogonal click-chemistry — to investigate the trafficking and metabolism of sphingolipids. This approach offers high chemoselectivity and fast processing of broad sample types (40–42), making it an appealing tool to answer questions in host-microbe interactions. For example, copper (I) catalyzed azides alkynes cycloadditions are regarded as the archetype of click-chemistry (43), and are especially useful for our application.

Therefore, we developed an integrated methodology that permits comprehensive identification and characterization of the fates of dietary sphingolipids delivered to the gut microbiome. In this study, omega-alkynyl sphinganine (sphinganine alkyne – SAA), a sphinganine (SA) surrogate containing an alkyne group, was introduced to mice by oral administration (Click). After biological processing, fluorescent dyes were then conjugated to SAA-containing metabolites by click-chemistry. These fluorescent features permit the use of fluorescence-activated cell sorting (FACS) to detect and isolate microbes with alkyne-tagged sphingolipid metabolites in a high-throughput manner (Sort), which further enables downstream applications such as 16S rRNA gene sequencing (Seq). Finally, by leveraging the specific qualities of the alkyne, SAA-derived metabolites are revealed through the use of liquid chromatography coupled to high-resolution mass spectrometry (LC-HRMS) (Spec).

Application of Click-Sort-Seq-Spec (ClickSSS) (Fig. 1) revealed a set of microbes that take up dietary sphingolipids. This included the sphingolipid-producing *Bacteroides* and beneficial microbes such as *Bifidobacterium* that do not produce sphingolipids. Comparative metabolomics revealed the diet-dependent sphingolipidome of both *Bacteroides thetaiotaomicaron (B. theta)* and *Bifidobacterium longum* subsp. *infantis (B. longum)*. Our analysis suggests that diets rich in sphingolipids can influence the composition of the microbiome with implications for supporting the colonization of beneficial microbes. Moreover, ClickSSS can be applied to a variety of diet-microbiome systems to uncover the mechanism of nutrient interactions with the gut microbial metabolome.

**Fig. 1. -.**
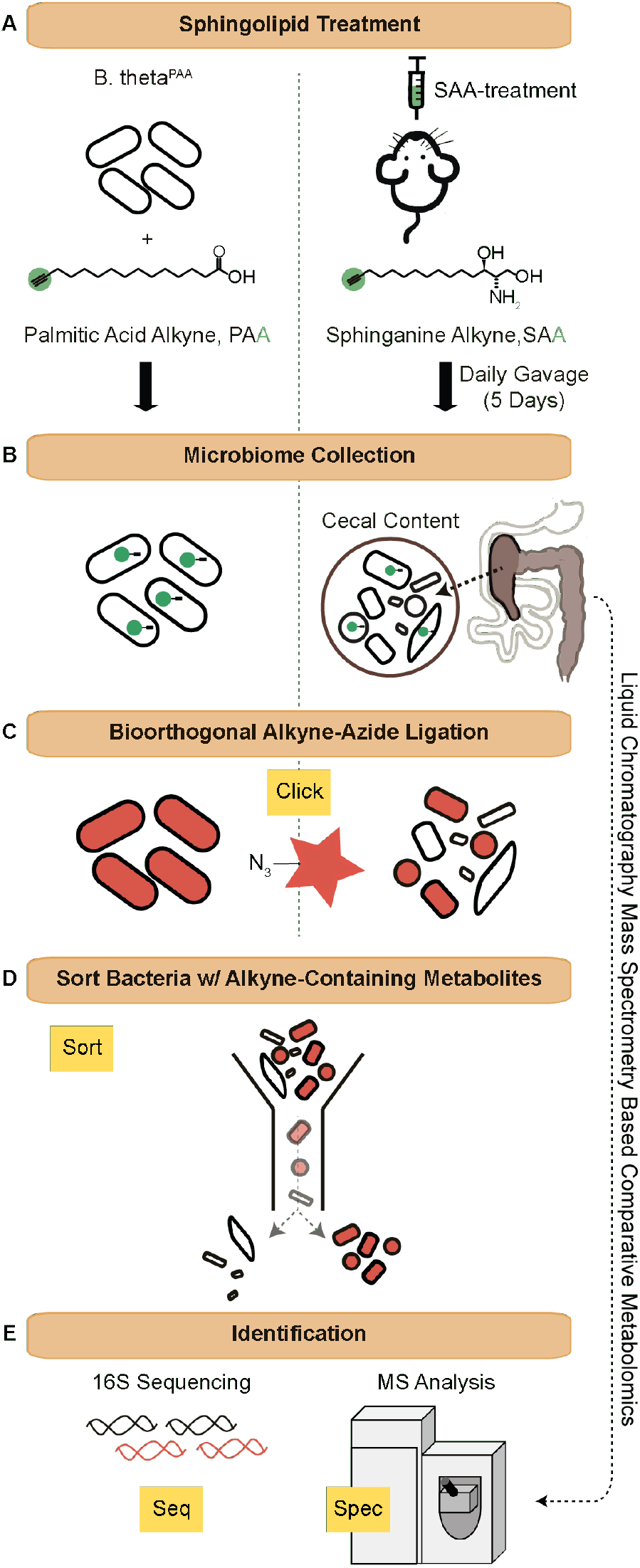
Click-Sort-Seq-Spec (ClickSSS) workflow. (A) An alkyne containing lipid was introduced to either the *Bacteroides thetaiotamicron* (*B. theta*) cultures (palmitic acid alkyne, PAA) or to mice by oral gavage (sphinganine alkyne, SAA). (B) *B. theta* labeled with PAA (*B. theta*^PAA^) was collected after a 24-hour incubation and the cecal content from mice was harvested after oral administration of SAA for five consecutive days. (C) Sphingolipid-interacting bacteria were fluorescently detected with Alexa Fluor 647 azide using click chemistry. (D) Sphingolipid-interacting versus non-interacting bacteria were separated using fluorescence-activated cell sorting (FACS). (E) Species composition of interacting versus noninteracting bacteria was determined by 16S sequencing. The metabolic consequences of PAA and SAA exposure were determined by differential metabolomic analysis.

## Materials and Methods

### General Information

All buffer, reagents, media, and instrumentation were utilized sterile. All solvents used for metabolomics were purchased from Fisher Scientific as HPLC grade.

### Bacterial culturing

*Bacteroides thetaiotaomicron* strain VPI 5482 (*B. theta*) was cultured in modified minimal medium (44) consisting of 13.6 g KH_2_PO_4_, 0.875g NaCl, 1.125 g (NH_4_)2SO_4_, 5 g glucose, (pH to 7.2 with concentrated NaOH), 1 mL hemin solution (500 mg dissolved in 10 mL of 1M NaOH then diluted to final volume of 500 mL with water), 1 mL MgCl_2_ (0.1M in water), 1 mL FeSO_4_x7H_2_O (1 mg per 10 mL of water), 1 mL vitamin K_3_ (1 mg/mL in absolute ethanol), 1 mL CaCl_2_ (0.8% w/v), 250 μL vitamin B_12_ solution (0.02 mg/mL), and 5g L-cysteine hydrochloride anhydrous. *Bifidobacterium longum* subsp. *infantis (B. longum)* ATCC 15697 was stored in deMann, Rogosa, and Sharpe (MRS) broth (Difco, Sparks, MD, USA) containing 60% (v/v) glycerol at −80°C. An MRS agar plate was streaked from the frozen stock and incubated at 37°C under anaerobic conditions overnight. One colony from the MRS agar plate was propagated into MRS broth and incubated at 37°C overnight. Culture broths were prepared in an anaerobic chamber (Coy Laboratory Products Inc., Grass Lake, MI) maintained with a gas mixture of 20% CO_2_, 5% H_2_, and 75% N_2_, then aliquoted into gas-tight jars. Incubation was performed at 37°C. Cultures were grown to an OD_600_ of 0.4-0.5 prior to use.

### *In vitro* cultures of *B. theta* and *B. longum* treated with alkyne lipids

*B. theta* was cultured with either 25 μM palmitic acid alkyne (PAA) or vehicle alone (ethanol) anaerobically at 37°C. For metabolic profiling experiments, *B. theta* and *B. longum* were incubated with or without 25 μM sphinganine alkyne (SAA) (Click Chemistry Tools, Scottsdale, AZ), under anaerobic conditions at 37°C. After 24 hours of incubation, the cultures were harvested by centrifugation at 18,000 × g for 10 min at room temperature. Cell pellets were washed with phosphate-buffered saline (PBS) and then washed three times with 1% BSA/PBS. Washed cell pellets were either fixed with 4% paraformaldehyde (PFA) in PBS for labeling with fluorophore Alexa Fluor 647 azide (AF647-azide, Invitrogen, Carlsbad, CA) or flash frozen with liquid nitrogen and stored at −80°C until additional processing for LC-HRMS analysis.

### *In vivo* dietary sphingolipids uptake

Murine experimental procedures were approved by the Cornell University Institutional Animal Care and Use Committee (IACUC) protocol #2010-0065. Thirty-two female 5-week-old Swiss Webster mice were purchased from Taconic Biosciences and subjected to vivarium habituation for one day after delivery. Mice were randomly assigned into four treatment groups. Each treatment consisted of eight mice housed 4 to a cage. Mice were all fed a standard sterilized breeder diet (LabDiet 5021, St. Louis, MO). Dietary sphinganine was introduced by oral gavage in 100 uL of 50/50 dimethyl sulfoxide (DMSO) and PBS with sphinganine (SA) or SAA at 5 mg/kg of body weight. Mice were gavaged daily with either treatment, vehicle control or no gavage for 5 days. Fecal pellets were collected from each cage at the time of sacrifice, temporarily moved to ice and then stored at −80°C for subsequent analyses. Five hours after the final gavage, all the mice were exsanguinated by decapitation. Cecal contents were collected and snap-frozen, and stored at −80°C until further processing.

### Isolation and fixation of bacterial cells from cecal content samples

Cecal contents were thawed at room temperature for 3-5 minutes. To separate microbial aggregates and individual cells from fibrous debris, cecal contents were weighed and diluted 1:10 with ice-cold 1X PBS in a 1.5 mL tube. Samples were then vortexed for 5 min, and then mildly sonicated for 20s total on time, with alternating 2 second on pulses, and 2 second off pulses, at 3-6W (Qsonica Ultrasonic Processor, Model Q700, with a water bath adaptor, Model 431C2). Between pulsing intervals, the samples were chilled on ice for 20s. Samples were centrifuged for at room temperature for 2 min at 200 × g and supernatants were transferred to a fresh tube. To maximize recovery, the remaining pellets were subjected to a second round of isolation. The supernatants were pooled and centrifuged at 18,000 × g for 10 min. The bacterial pellets were then washed three times with ice-cold 1% bovine serum albumin (BSA) in PBS. After the final centrifugation, the supernatants were discarded, and the bacterial cell pellet was resuspended in 4% PFA in PBS for 10 min. Cells were then gently washed with 1% BSA/PBS and isotonic Triton X buffer (0.5% Triton X-100 in PBS) was added for permeabilization for 10 min at room temperature.

### Cu(I)-catalyzed azide-alkyne cycloaddition staining

Bacterial cells containing PAA or SAA derivatives were labeled with AF647-azide using freshly prepared click-reaction cocktail prepared according to manufacturer’s instructions for the Click-&-Go™ Click Chemistry Reaction Buffer Kit (Click Chemical Tools, Scottsdale, AZ) at a final fluorophore concentration of 5 μM for 30 min at room temperature. Cell pellets were washed 5 times and resuspended with 1% BSA/PBS to remove any non-specific fluorescent signals before flow cytometry analysis.

### Fluorescence-activated cell sorting to identify SAA containing bacteria

A BD Biosciences Melody Sorter (BD Biosciences, San Jose, CA) was sterilized by subsequent runs of 70% ethanol and autoclaved distilled water. The instrument was kept under sterile conditions during analysis and sorting by use of sheath fluid prepared using sterile 1X PBS. Samples were passed through a 35μm nylon mesh cell strainer (Corning, Amsterdam, the Netherlands). The AF647-azide dye was excited using a 640-nm red laser and fluorescence was captured with a 660-nm/ 20-nm filter. The initial gate was developed and drawn on the AF647 positive events using the *B. theta*^PAA^ sample which was confirmed to contain alkyne-bearing metabolites, including sphinganine alkyne (SAA), by both fluorescence imaging and targeted metabolomics (Fig. 2A and 2B) as described below. To remove potential cell debris (debris exclusion) and false-positive signals detected from clumped cells (doublet exclusion) in the cecal content sample, the gating was further optimized using combinations of forward scatter (FSC) and side scatter (SCC) parameters (Fig. 4). To ensure that debris and doublet exclusion steps did not affect the initial AF647 positive gate or exclude bacterial cells, the *B. theta*^PAA^ sample was reanalyzed according to the modified sort parameters of the cecal samples (Fig. 4A). The cecal content sample from mice in the vehicle control group (untreated) was subjected to the above gating steps and further used to draw the AF647 negative gate. Finally, AF647-positive and AF647-negative gates were both applied to cecal content from mice in the SAA group (SAA treated) after the debris and noise removal gating steps.

**Fig. 2 -.**
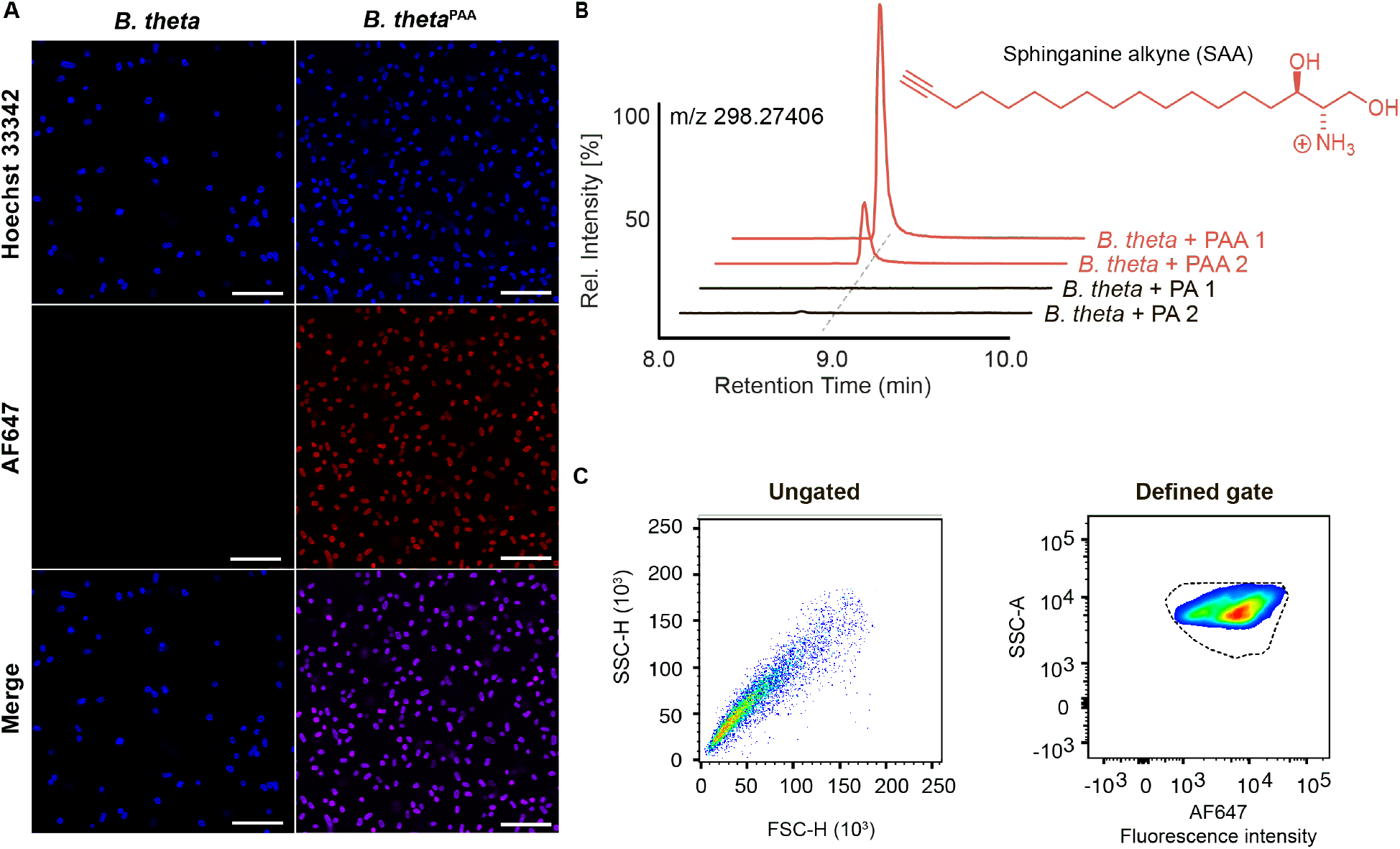
Development of sorting parameters using *B. theta*. (A) Uptake and assimilation of PAA by *B*. *theta* was visualized by fluorescence microscopy in the red channel after incubation of cultures with Alexa Fluor 647 (AF647) azide. DNA was stained using Hoechst 33342 and visualized in the blue channel (Scale bar: 20 μm). (B) Ion chromatogram and chemical structure of sphinganine alkyne (SAA) detected in *B. theta*^PAA^. PA = palmitic acid, PAA = palmitic acid alkyne, FSC-H = forward scatter height, SSC-H = side scatter height. (C) Dots and density plots showed the distribution of *B. theta*^PAA^ detected by AF647-azide before (ungated) and after (defined gate) defining gates for AF647 positive and negative populations by flow cytometry.

After establishing this gating strategy, 60,000 events in each gate with an AF647 signal greater than 10^3^ fluorescent intensity (AF647-positive) or 0 to 10^−2^ fluorescent intensity (AF647-negative) were captured and segregated in 2 separate 5 ml round-bottom tubes (BD Biosciences, San Jose, CA) containing 1% BSA/PBS. A second round of sorting was performed on the AF647-positive fraction with the parameters above. After sorting, cells were transferred to 1.5 mL tubes and centrifuged at 18,000 × g for 10 min at 4°C. The supernatant was discarded, and the sorted cells were counterstained with 1 μg/mL Hoechst 33342 (Invitrogen, Carlsbad, CA), then washed and resuspended with 1% BSA/PBS for imaging or 16S sequencing.

### Optical microscopy

Resuspended cells were mounted onto glass slides with Vectashield Vibrance mounting media (Vector Labs, Burlingame, CA) and analyzed using a Leica DM500 fluorescence microscope (Leica, Buffalo Grove, IL) or a Zeiss LSM 880 (Carl Zeiss, Jena, Germany) inverted confocal microscope, all the images were analyzed using Fiji Image J software (45). Additionally, the supernatant from each microbial isolation from cecal contents was imaged for confirmation of the absence of bacteria with Hoechst 33342 (1 μg/mL) and imaged using a fluorescence microscope. Samples were imaged by using two color filter cubes—UV (359 nm/461 nm) for Hoechst to localize cells, and Cy5 (650 nm/670 nm) for AF647 to detect the presence of alkyne-containing metabolites.

### DNA extraction

DNA was isolated according to (46). Specifically, the bacterial pellets were resuspended in TE buffer comprising of 10 mM Tris, 1 mM EDTA (pH 8.0), and processed with 3 consecutive freeze-thaw cycles in liquid nitrogen alternated with a 95°C incubation. After adding 10% 1M sodium dodecyl sulphate (SDS) and 10 μL of 8 U/mL proteinase K (NEB, #P8107S, Ipswich, MA), the suspension was then incubated at 56°C for 4 hours. 0.5 M NaCl and 65°C pre-heated 0.5 M cetyltrimethyl ammonium bromide were added and the well-mixed samples were then incubated at 65°C for 10 min to lyse the cells. After centrifugation at 18,000 × g for 5 min at 4°C, supernatants were transferred to fresh 2 mL microcentrifuge tubes, and 900 μL of phenol/chloroform/isoamyl alcohol (25:24:1, pH = 6.7) was added to each extraction. After mixing thoroughly, samples were incubated at room temperature for 10 min. To perform phase separation, samples were first centrifuged at 18,000 × g for 10 min at 4°C. Then the upper aqueous phase was collected and re-extracted with a further addition of 900 μL phenol/chloroform/isoamyl alcohol. Solutions were centrifuged at 18,000 × g for 10 min at 4°C, and the upper aqueous phases were transferred to fresh 2 mL microcentrifuge tubes. The final extraction was performed with the addition of 900 μL chloroform/isoamyl alcohol (24:1), and then subject to centrifugation at 18,000 × g for 10 min at 4°C for layer separation. The upper aqueous phase was transferred to a fresh 2 mL microcentrifuge tube. To precipitate a higher yield of DNA, 450 μL of isopropanol was added, and samples were incubated in a −20°C freezer overnight. Finally, DNA was collected by centrifugation at 18,000 × g for 30 min at 4°C. The supernatant was discarded and the pellets were air-dried and dissolved in TE buffer to a final volume of 30 μL.

### 16S rRNA gene sequencing

Amplicon libraries were created by PCR using universal bacterial primers targeting the V4 region of the 16S rRNA gene with primers 515F (5’-AATGATACGGCGACCACCGAGATCTACACTATGGTAATTGTGTGCCAGCMGCCGCGGTAA-3’) and 806R (5’-CAAGCAGAAGACGGCATACGAGAT XXXXXXXXXXXX AGTCAGTCAG CC GGACTACHVGGGTWTCTAAT-3’) (47). Barcoded forward and nonbarcoded reverse primers were used with Taq DNA polymerase Master Mix (TONBO biosciences, CA) according to the manufacturer’s directions. Samples were amplified in duplicate with the following thermocycler protocol: hold at 94°C for 3 min; 25 cycles of 94°C for 45 s, 50°C for 1 min, 72°C for 1.5 min; and hold at 72°C for 10 min, and the duplicate final amplified products were pooled. 16S rRNA gene amplicons were cleaned with Mag-Bind^®^ RxnPure Plus (Omega Bio-tek, Inc., GA). Samples were mixed together in equimolar amounts prior to sequencing on Illumina’s MiSeq platform. Sequence data was processed in QIIME 2 (Quantitative Insights Into Microbial Ecology 2) (48). Libraries were demultiplexed, raw sequence pairs were joined, and the quality was trimmed based on an average quality score of 30. Sequences were clustered into *de novo* OTUs with 97% similarity using the Greengenes database (version 13.8) (49). The negative controls run with each MiSeq set for determining potential contaminants in PCR reagents and procedure were analyzed and removed (50).

### Preparation of cecal contents for analysis by LC-HRMS

Frozen cecal contents were thawed at room temperature for 3-5 min. 500 μL of PBS was added and the slurry was vortexed for 1 min, followed by centrifugation for 2 min at 200 × g at room temperature. The supernatant was collected, a second round of PBS was added, and the process repeated. The supernatants were pooled and filtered through a 35 μm nylon mesh cell strainer (Corning, Amsterdam, the Netherlands). After centrifugation at 18,000 × g for 10 min at 4°C, the supernatant was discarded. The residual bacterial pellets were homogenized in a 2 mL microtube containing 1 mm zirconium beads (OPS diagnostics, NJ) and 500 μL of PBS for 3 min using a mini-BeadBeater (BioSpec products, Bartlesville, OK). Afterward, the lysate was placed on ice for 5 min to cool down and 10 μL of the lysate was added to 90 μL RIPA buffer (Thermo Scientific, Waltham, MA) for determination of protein concentration. The Lowry protein assay (kit from BioRad Laboratories, Hercules, CA) was used to standardize protein concentrations. 1000 μg of protein per bacterial pellet sample was transferred to a fresh 1.5 mL tube (USA Scientific, Ocala, FL). Lysates were then frozen with liquid nitrogen and lyophilized to dryness.

### Lipid extraction for LC-HRMS analysis

1 mL of methanol was added to the dried lysate and sonicated for 1 min, on/off cycles of two seconds on, three seconds off, at 100% power on a Qsonica Ultrasonic Processor. The samples were then placed on an end over end rotator and metabolites were extracted overnight. Samples were then centrifuged at 18,000 × g at 4°C for 30 min. The clarified supernatant was collected and transferred to a fresh 1.5 mL centrifuge tube. The collected extracts were evaporated to dryness with a SpeedVac vacuum concentrator (Thermo Fisher Scientific, Waltham, MA) and then reconstituted in 200 μL of methanol. Samples were sonicated again (*vide supra*) and then centrifuged at 18,000 × g at 4°C for 30 min. 150 μL of clarified concentrated extracted metabolome was transferred to an HPLC vial utilizing an insert (Thermo Scientific, Waltham, MA) and stored at 4 °C until LC-HRMS analysis.

### Mass Spectrometric Analysis

High resolution LC-MS (LC-HRMS) analysis was performed on a Thermo Fisher Scientific Vanquish Horizon UHPLC System coupled with a Thermo Q Exactive HF hybrid quadrupole-orbitrap high-resolution mass spectrometer equipped with a HESI ion source. 3 μL of extract was injected and separated using a water-acetonitrile gradient on a Kinetex EVO C18 column (150 mm × 2.1 mm, particle size 1.7 μm, part number: 00F-4726-AN) maintained at 40°C. Solvent A: 0.1% formic acid in water; Solvent B: 0.1% formic acid in acetonitrile. A/B gradient started at 10% B for 3 min after injection and increased linearly to 100% B at 20 min and held at 100% B for 10 min, using a flow rate 0.5 mL/min. Mass spectrometer parameters: spray voltage 3.5 kV for positive mode and 3.0 kV for negative mode, capillary temperature 380°C, prober heater temperature 400°C; sheath, auxiliary, and spare gas 60, 20, and 1, respectively; S-lens RF level 50, resolution 240,000 at m/z 200, AGC target 3× 10^6^. Each sample was analyzed in positive and negative modes with m/z range 100 to 1200.

### Untargeted Metabolomic Analysis

RAW files generated from HPLC-HRMS acquisitions were converted to mzXML files utilizing MSconvertGUI software (proteowizard.sourceforge.net). Differential molecular features were determined by subjecting mzXML files to Metaboseek Software version 0.9.6 (metaboseek.com) utilizing the xcms package (51). Differential features were filtered using the minFoldOverCtrl, minInt, and Fast_Peak_Quality filters, and then subjected to manual curation to remove adducts and isotopes. The curated features were assigned molecular formulas and then subjected to tandem mass spectrometry (MS2). MS2 fragments were also assigned molecular formulas and sphingolipid structures were inferred.

## Results

### ClickSSS can identify microbes that uptake dietary lipids

To understand how dietary sphingolipids interact with the microbiome, we developed a strategy to determine the fate of orally introduced alkyne tagged sphingolipids. This involved the isolation of microbial cells from the cecal content of mice that were orally gavaged with sphinganine alkyne (SAA) for 5-days. The alkyne functional group allows detection of alkyne containing molecules through the copper catalyzed cycloaddition of an azide conjugated detection reagent – a type of click chemistry. These bacterial pellets were then incubated with an azide conjugated fluorophore (Alexa Fluor 647 azide, AF647-azide) so that microbial cells that assimilated dietary SAA could be identified by either microscopy or flow cytometry (Fig. 1). To overcome the challenges of sorting mixed populations of microbial cells, we initiated method development with laboratory-friendly *Bacteroides thetaiotamicron* (*B. theta*), a gut commensal bacteria also known to produce sphingolipids. The first committed step in the biosynthesis of sphingolipids relies on the conjugation of a long chain fatty acid, such as palmitic acid, to serine via the enzyme serine palmitoyltransferase (SPT). Previous work showed that *B. theta* assimilates the alkyne containing fatty acid, palmitic acid alkyne (PAA) (52), which provided a platform for us to sort PAA exposed versus unexposed cultures. We therefore fed PAA to *B. theta* to enable the production of alkyne-bearing lipids. PAA was added to axenic cultures of *B. theta* for one day and the cells were collected via centrifugation. Cells were then washed, fixed, permeabilized, and then covalently ligated (“clicked”) to AF647-azide (Fig. 1A-C).

Fluorescence imaging showed overlapped Hoechst and AF647 signals in the PAA-treated samples that were absent from untreated samples, indicating successful integration of the alkyne label into *B. theta* metabolites (Fig. 2A). This was further validated via targeted metabolomics showing that PAA-treated *B. theta* gives rise to alkyne-bearing sphinganine (SAA), the expected product of SPT ligation of PAA and serine (Fig. 2B). We then applied both the PAA-treated and untreated bacteria to fluorescence activated cell sorting (FACS), of which we were able to appropriately determine gating of AF647-positive cells. The density plot obtained for PAA-treated cells (*B. theta*^PAA^) showed that detectable cells presented a narrow fluorescent intensity distribution, indicating that PAA integration into the *B. theta* metabolome was rather effective (Fig. 2C). Thereafter we drew an initial gate based on the contour of *B. theta*^PAA^ sample (Fig. 2C, defined gate).

### Sphinganine alkyne (SAA) is assimilated by select gut microbiota

To determine the presence and identity of dietary sphingolipid interacting bacteria, adult female mice were gavaged with ~75 mg/day of SAA for five consecutive days. The cecal contents were collected and individual cells were detached from neighboring cells and matrix particulates by mild sonication and centrifugation, which provided preferential cell recovery for downstream analyses (53, 54). Click chemistry of AF647-azide was applied to both SAA-treated and -untreated cecal microbial consortia. To ensure that extracted cells showed detectable AF647 signal prior to physical separation, non-destructive fluorescence microscopy was utilized. Our results showed that only a portion of the bacteria were positive for the AF647 signal against the Hoechst 33342 signal (Fig. 3). This result suggested that a select fraction of the microbiome was able to uptake dietary sphinganine while the remaining fraction did not appear to interact with this introduced metabolite.

**Fig. 3.**
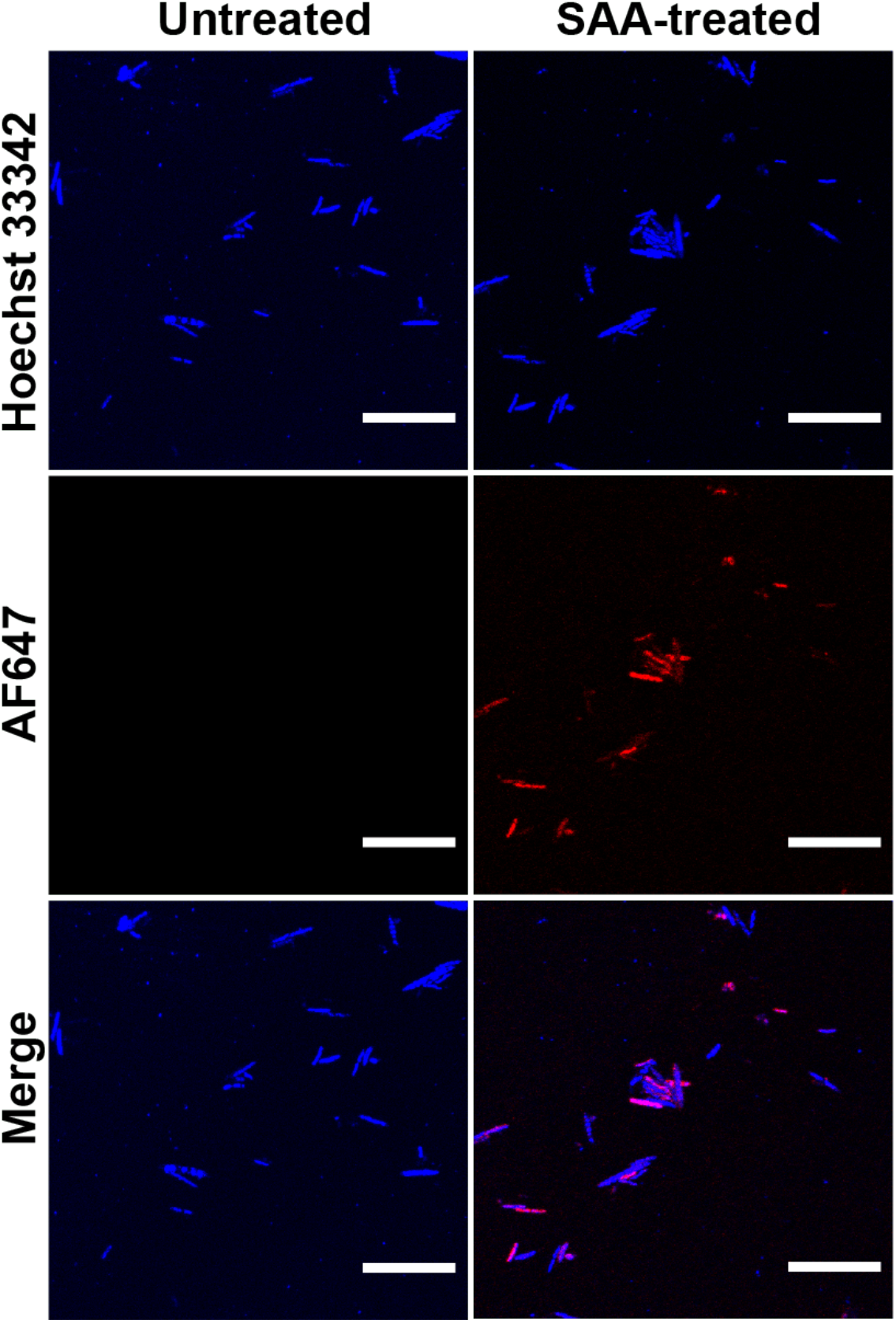
Selective assimilation of dietary sphinganine by the gut microbiome. Confocal images of bacteria extracted from the cecal content of mice orally exposed to sphinganine alkyne (SAA-treated) versus the vehicle control group (untreated). SAA interacting microbes are red (AF647 azide) while a general DNA stain marks all bacteria in blue (Hoechst 33342). (Scale bar: 20 μm).

### 16S sequencing of sorted microbiome samples revealed the identity of sphingolipid-interacting bacteria

To reveal the identity of SAA interacting bacteria, we sought to separate AF647 positive and AF647 negative microbes by FACS (Fig. 1D). After bacteria were isolated from bulk cecal content and then conjugated to AF647 azide, the bacterial cells were then subjected to FACS. Cells were first gated for size and shape using light scattering. Initial observations showed that relative to *B. theta*^PAA^ samples, the distribution of the AF647-treated cecal contents were more dispersed under side scatter height (SSC-H) and forward scatter height (FSC-H) assessment (Fig. 4A, ungated), suggesting background noise from the cecal matrix. In order to improve the signal-to-noise ratio, the SAA-untreated cecal content samples were used to determine the background noise (Fig. S1A, overlapped area marked as yellow), and two additional gates were applied to exclude debris and doublets which contribute to false-positive signals (Fig. 4A). Further evaluation of the fluorescence cytograms established two distinct populations, identified as either high or low fluorescence intensity. To ensure that the revised gates set for removal of debris and doublets would not alter the outcome defined by our initial gates established for *B. theta*^PAA^, *B. theta*^PAA^ were then subjected to the revised method. Our optimized method showed no significant exclusion of the bacterial cells of interest (Fig. 4A). Therefore, the gate developed from AF647 positive *B. theta*^PAA^ was applied to all AF647 positive microbial populations. From this point forward, we defined the cells eluting from this gate as AF647-positive and the population shown in the low fluorescence gate as AF647-negative. The negative gate was established by the SAA-untreated cecal sample to control for background fluorescence (Fig. 4A, Fig. S1B defined gate). We then sorted ~60,000 events from cecal contents of SAA treated mice for each of these two gated populations. To ensure removal of dimeric or anomalous features, AF647-positive cells underwent an additional round of FACS. Purity of the sort was further confirmed by fluorescence microscopy (Fig. 4B).

**Fig. 4.**
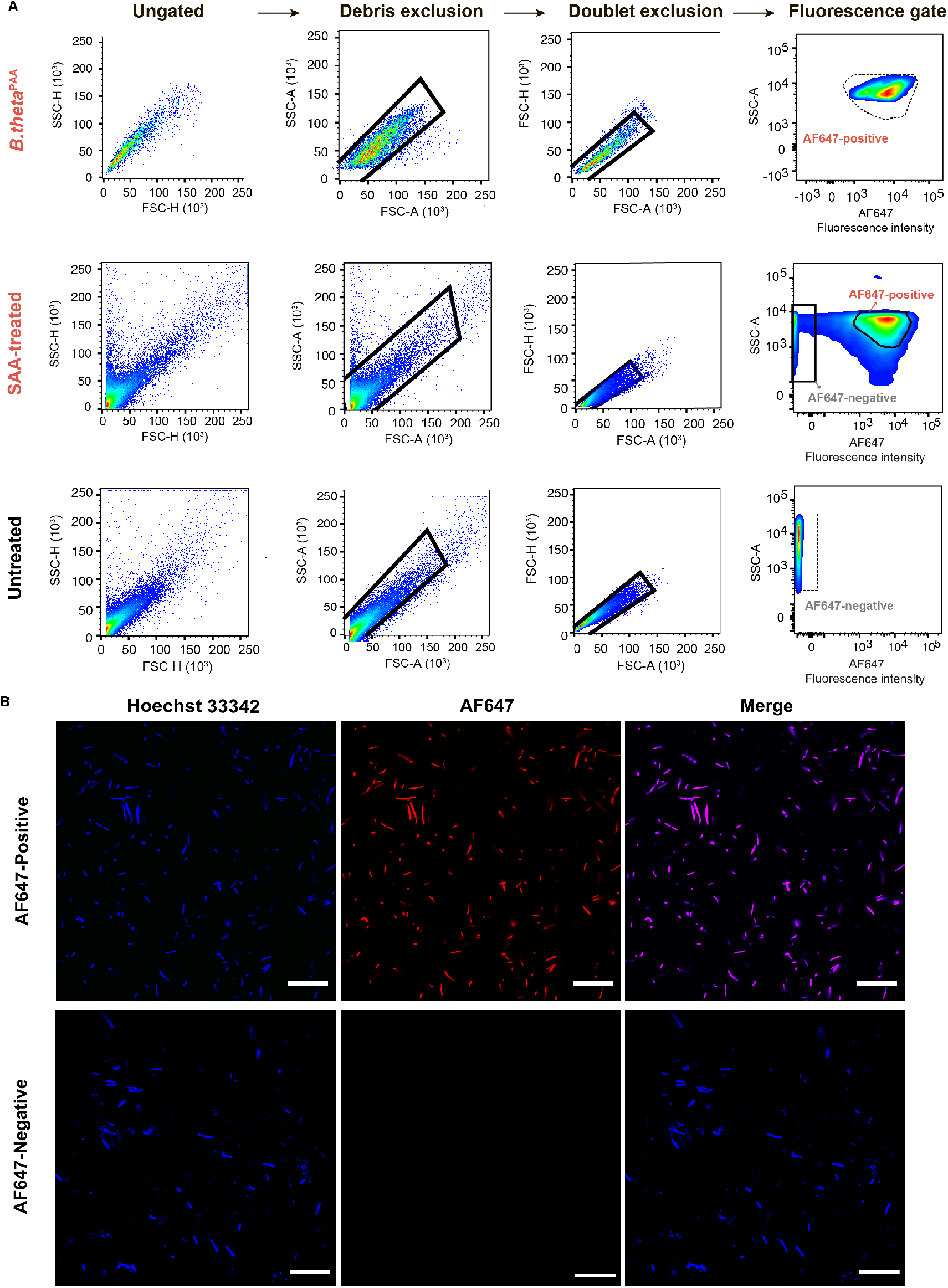
Sphingolipid-interacting and non-interacting bacteria can be separated using FACS. (A) Representative density plots showing gates used to sort AF647 positive versus AF647 negative populations. The successive steps are outlined on the top of the plots and the experimental group is identified on the left side of each row. The solid line indicates the defined gate for the target fractions and the dashed line indicates the derivation of those two gates from the *B. theta*^PAA^ and SAA-untreated samples. (B) Representative fluorescence microscopy images confirming the purity of the two bacterial populations after FACS. SAA interacting microbes are red (AF647 azide) while a general DNA stain marks all bacteria in blue (Hoechst 33342). (Scale bar: 20 μm).

To determine the identity of the AF647-positive microbes, and therefore bacteria containing SAA or SAA-derived metabolites, we sequenced the 16S rRNA gene (16S sequencing) of the cells obtained by FACS (Fig. 1E). Biological duplicates of ~60,000 events retrieved by FACS from two mice were used for 16S sequencing. Operational taxonomic units (OTUs) at the genus level in the AF647 positive sample were ranked in descending order corresponding to their relative abundance (Fig. 5A). Interestingly, the AF647-positive bacteria, was dominantly composed of *Bacteroides* spp. (99%). The remainder of AF647-positive bacteria was composed of, in the order of prevalence, *Prevotella* spp.*, Lactobacillus* spp., and *Bifidobacterium* (Fig. 5B). *Bacteroides* spp. and *Prevotella* spp. are known sphingolipid producers, suggesting that sphingolipid-producing (SP) gut bacteria play a major role in processing dietary sphinganine. Additionally, there is currently no research demonstrating the ability of either *Bifidobacterium* or *Lactobacillus* to generate sphingolipids *de novo*, suggesting a yet unexplored metabolic link between sphingolipids and non-sphingolipid-producing (NSP) bacteria. To this end, we wished to better understand the metabolic consequences of dietary SA uptake by the microbiome and explore the cecal microbial metabolomes of SAA-treated mice.

**Fig. 5.**
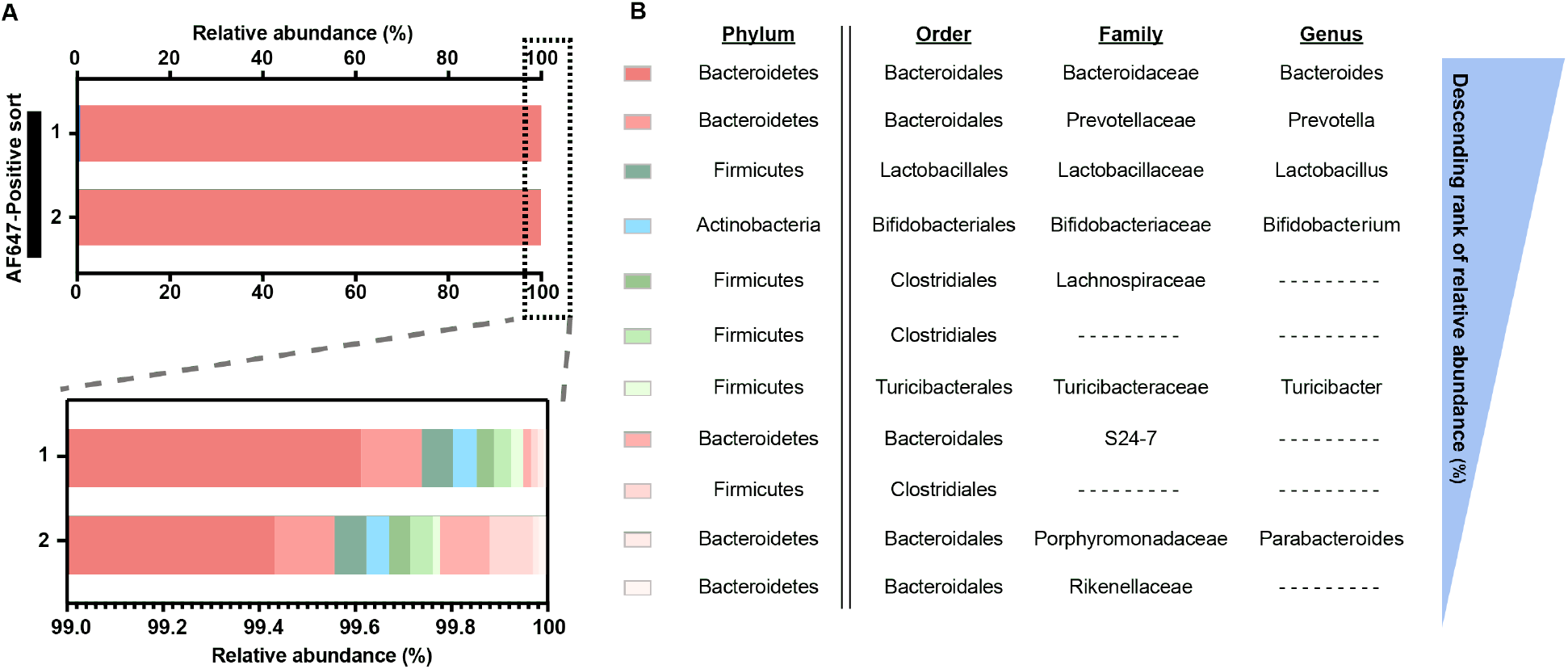
Identity of sphingolipid-interacting and non-interacting gut microbes. (A) Bacterial DNA from FACS sorted AF647-positive cecal content samples was used to determine microbiome composition using 16S sequencing. The bar graph shows the relative abundance (%) of each operational taxonomic unit (OTU) at the genus level in the total AF647 positive fraction. (B) The focused bar chart shows the microbial diversity of the minor fraction (1%) of the sphingolipid-interacting fraction. (C) OTUs are ranked in descending order of the relative abundance in the AF647 positive fraction.

### SAA is metabolized by the cecal microbial consortium

To determine if the gut microbial consortium was metabolizing SAA, we carried out comparative metabolomics via liquid chromatography coupled to high resolution mass spectrometry (LC-HRMS)(Fig. 1C,E). Conveniently, the unligated alkyne tag serves as a marker which allows us to trace the metabolic flux and fate of SAA. Inspection of SAA-treated cecal consortium metabolomes revealed the production of dihydroceramides (DHCeramide) consistent with N-acyl addition of fatty acids known to be produced by sphingolipid-producing bacteria. Differential metabolic features were observed between 17 and 20 minutes along the chromatograph corresponding to DHCeramides bearing fatty acyl side chains containing between 15- to 22-carbons (C15 to C22) with various oxidation statuses (Fig. 6, Fig. S2).

**Fig. 6.**
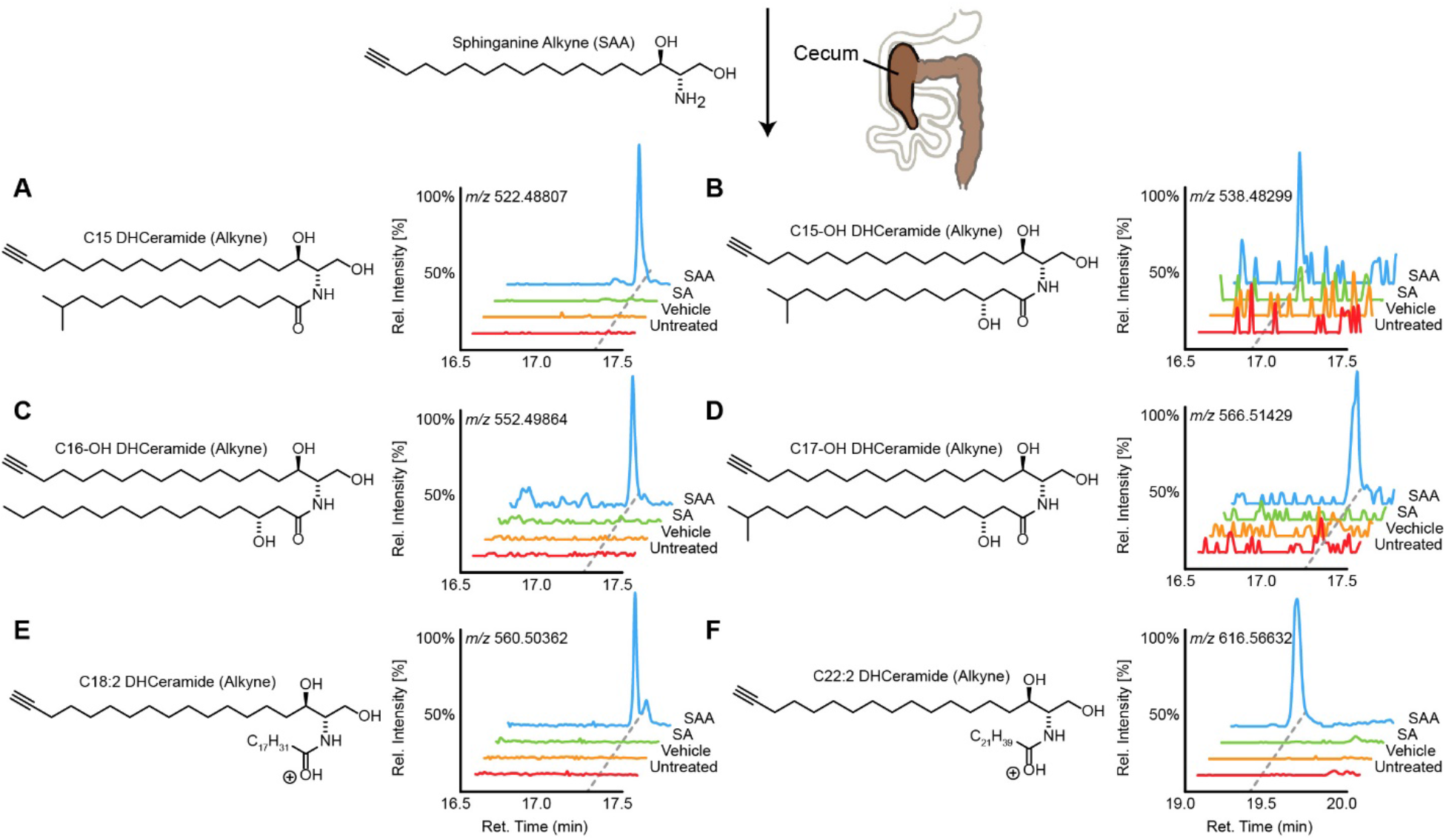
The gut microbiome transforms dietary sphinganine into dihydrocermides (DHCeramide). Representative structures and high-resolution mass spectrometry ion chromatograms of alkyne-bearing (A) C15-, (B) C15OH-, (C) C16OH-, (D) C17OH-, (E) C18:2-, and (F) C22:2-DHCerarmide from mice cecal microbial metabolomes orally treated with sphinganine alkyne (SAA, blue), but not detected in treatments of sphinganine (SA, green), vehicle, or no treatment (red). The x-axis represents retention time along a reverse phase column and the y-axis represents relative intensity normalized to the largest peak in the time window.

### SAA is metabolized to long chain ceramides via *Bacteroides* and not *Bifidobacterium*

To determine if SAA-derived long chain ceramides were constructed solely by sphingolipid-producing bacteria (SP), we grew isolated cultures of either *B. theta* or *Bifidobacterium longum (B. longum)* with SAA, representing our SP and NSP bacteria, respectively. *In vitro* cultures of *B. theta* treated with SAA produced the corresponding C15-, C16-, and C17-ceramides found in cecal metabolomes (Fig. 7A). Inversely, SAA-treated *B. longum* cultures showed no detectable amounts of long chain ceramides. Interestingly, more comprehensive analysis of SAA-treated *B. longum* cultures showed the production of SAA-derived short chain fatty acyl (C1- to C4-) ceramides, including N-acylation of common fermentation metabolites pyruvate, lactate, and succinate (Fig. 7B, Fig. S3). Although these minor products in *B. longum* may explain their AF647-positive sorting, these short chain ceramides were not detected in cecal metabolomes.

**Fig. 7.**
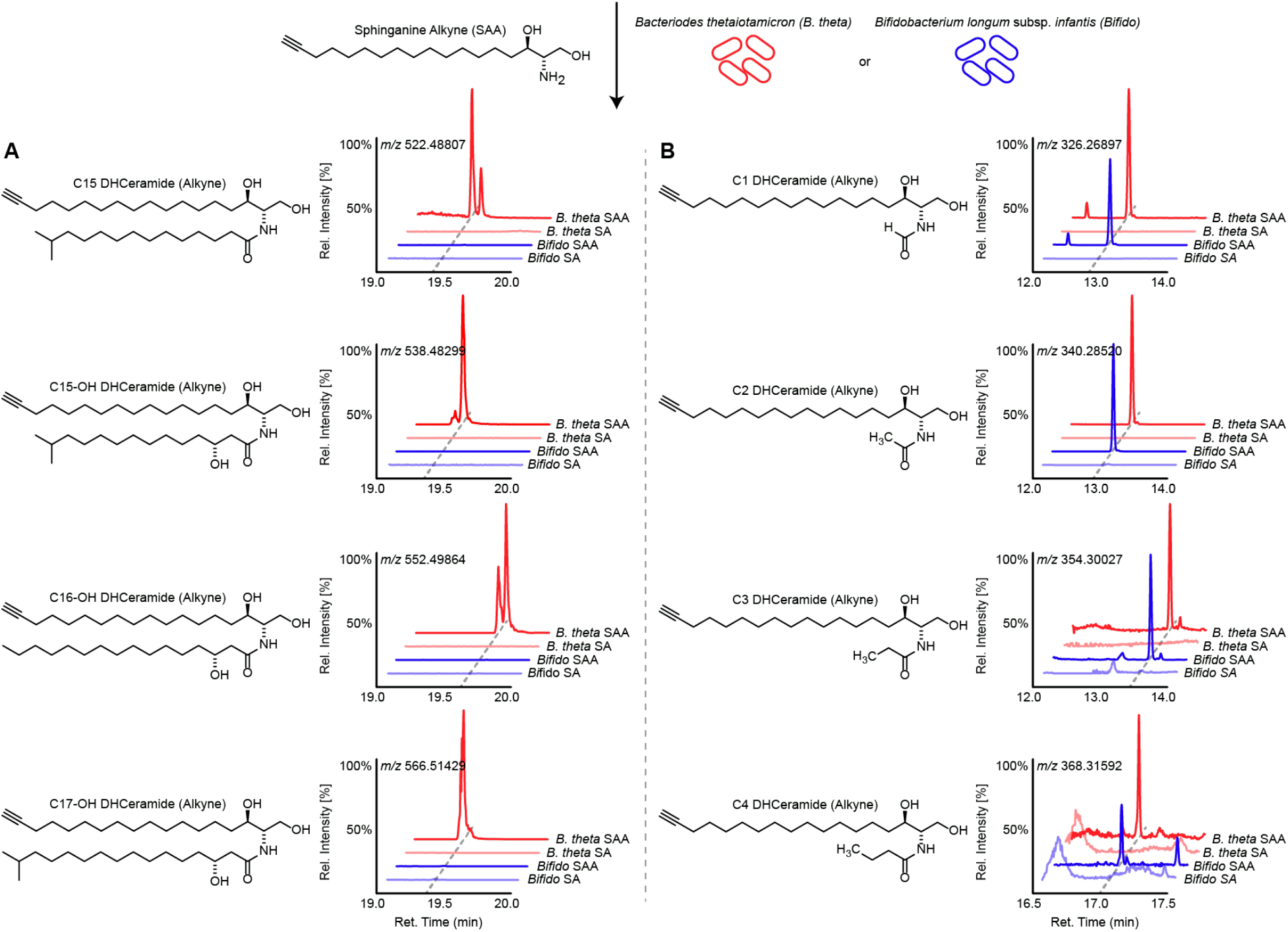
*B. theta* and *B. longum* make different dihydroceramides (DHCeramide) from exogenous sphinganine. Representative structures and high-resolution mass spectrometry ion chromatograms of metabolomes from *in vitro* cultures of *Bacteroides thetaiotamicron* (*B. theta*, red) and *Bifidobacterium longum* subsp. *infantis* (*B. longum*, blue) treated with either sphinganine alkyne (SAA, bold color) or sphinganine (SA, light color) representing the detection of SAA-derived alkyne bearing (A) long chain C15-17, and (B) short chain C1-4 DHCeramide. The x-axis represents retention time along a reverse phase column and the y-axis represents relative intensity normalized to the largest peak in the time window.

In addition, *B. theta* cultures produced significant levels of dihydroceramide phosphoethanolamides (Cer-PE), which could not be detected in cecal metabolomes (Fig. S4). Furthermore, longer chain (C18- to C22-) ceramides noted in the cecal metabolomes were not produced by isolated cultures of *B. theta* (Fig. S5). Taken together, these data suggest a fascinating context specific link between SAA and various gut microbes, in particular *B. theta* and *B. longum*, and their respective observable lipidomes.

## Discussion

In this study, we introduce a click-chemistry based method, ClickSSS, to both probe and identify the incorporation of lipid alkynes in gut bacteria. Treating *in vitro* cultures of *B. theta* with PAA, followed by chemical ligation of AF647-azide, we developed a FACS-based method for sorting bacteria that assimilate the alkyne-tagged lipid versus those that do not. Then, by treating mice with SAA, the ClickSSS workflow determined over 99% of the SAA assimilating bacteria was *Bacteroides*. Interestingly, the second, albeit far less abundant taxa in the AF647 population, was *Prevotella*. These results suggested that dietary sphinganine was almost exclusively processed by known sphingolipid-producing bacteria. To this end, we performed comparative metabolomics on the cecal microbial community from mice which were treated with and without SAA. We noted the production of C15-, C16-, and C17-ceramides with N-acyl groups originating from acids bearing β-hydroxy groups, consistent with processing of SAA by *B. theta*. Interestingly, longer chain (C18- to C22-) ceramides were also differential in the cecal metabolome which were not observed with *in vitro* cultures of *B. theta*, suggesting context specific production of ceramides by *B. theta* or processing by another gut microbe. In addition, loss of detectable Cer-PE *in vivo* suggests rapid metabolism by either the host, the consortium, or context specific down-regulation of production by *B. theta*. Finally, we draw attention to *B. theta’s* ability to sequester SA/SAA and perhaps other lipids from its environment which suggests underlying mechanism which may confer competitive advantages for *B. theta* under certain dietary states.

Notably, two non-sphingolipid-producing (NSP) bacterial genera — *Bifidobacterium* and *Lactobacillus* — were also identified in the AF647-positive gate. Previous studies showed a positive correlation between milk sphingomyelin and the proliferation of *Bifidobacterium* in the mammalian gut (55–58). Moreover, the beneficial symbiont effects from the combination of *Lactobacillus casei* and *Bifidobacterium bifidum* and sphingomyelin were found in mice with colon cancer (59). In addition, *Bifidobacterium’s* well-established processing of human milk oligosaccharides (HMO) highlights the dietary effects of ceramide-attached HMOs, conventionally referred to as gangliosides. *Bifidobacterium* have been shown to digest HMOs from gangliosides, liberating free ceramides (60). This illustrates clear connections between *Bifidobacterium* and sphingolipid biology, however these links remain unexplored. Overall, these results draw attention to NSP sphingolipid interacting bacteria, from which either novel signaling paradigms or shared metabolism likely occur and encourages further analysis. Hence, our results show that ClickSSS may help unravel the yet uncharacterized relationship among sphingolipids and other beneficial microbes.

In conclusion, ClickSSS offers a simplistic workflow to track metabolic fingerprints and trace the contributors of sphingolipids metabolism in complex samples. Our method not only expands the lexicon of host-microbiome interaction, but also provides a complementarily protocol to study the *in situ* activity response of microbial communities to dietary sphingolipids, which can be applied to a variety of lipid investigations. In view of multiple channels-analysis in flow cytometry and the breadth of commercially available azide/alkyne-modified lipids, dual or perhaps multiple metabolic labels could be traced simultaneously via the ClickSSS protocol. Although our study connects sphingolipid metabolism and the gut microbiome, with an ever-expanding repertoire of commercially available alkynes and azides, our use of ClickSSS translates easily into the study of virtually any metabolite and with any FACS-amenable metabolic system.

## Data Availability Statement

16S rRNA gene sequencing data is available in its processed form in the supplementary materials with raw sequences deposited in the Sequence Read Archive under accession number PRJNA637116. All other data are contained in the main text or the supplementary materials.

## Acknowledgements

The authors would like to thank Rebecca Williams for her guidance with confocal imaging. We thank the Genomics Facility of the Biotechnology Resource Center at the Cornell University’s Institute of Biotechnology for their help with sequencing experiments and financial support through their seed grant program. Additional grant support for data collected on the *Zeiss LSM880 inverted confocal/multiphoton microscope (i880)* came from grants NYSTEM C029155 and NIH S10OD018516.

